# HInt: interaction-based homology discovery through accelerated genome-scale AlphaFold screening

**DOI:** 10.64898/2026.07.31.741991

**Authors:** Quentin Rouger, Pierre Paillard, Manon Thomet, Emma Touquet, Gwenaël Rabut, Emmanuel Giudice, Damien F. Meyer, Kévin Macé

**Affiliations:** Univ. Rennes, CNRS, Institut de Génétique et Développement de Rennes (IGDR) - UMR6290, T4-SECRET Team, 35000 Rennes, France; Univ. Rennes, CNRS, INSERM, Institut de Génétique et Développement de Rennes (IGDR) - UMR6290, U1305, Ubiquitin System Team, 35000 Rennes, France; CIRAD, UMR ASTRE, 97170 Petit-Bourg, Guadeloupe, France; ASTRE, Univ. Montpellier, CIRAD, INRAE, Montpellier, France

## Abstract

Identifying homologous proteins across deep evolutionary distances remains a major challenge because sequence and structural similarity progressively become undetectable over time. Although protein-protein interactions (PPIs) are often constrained by function and evolution, whether conserved interaction interfaces can provide an independent signal for homology detection has remained largely unexplored owing to the computational cost of proteome-scale interaction prediction. Here we introduce HInt (Homology by Interaction), an accelerated AlphaFold-based framework that enables practical proteome-scale PPI prediction through biologically informed pre-filtering and optimised high-throughput structure modelling. Using HInt, we establish interaction-based similarity as a third axis of homology detection. We show that conserved interaction interfaces reveal homologous relationships that remain inaccessible to conventional sequence- and structure-based approaches. Application of HInt to both prokaryotic and eukaryotic systems, together with experimental validation, uncovered a previously unrecognised VirB5 pilus-tip protein in the F-plasmid type IV secretion system and a previously unannotated F-box-like protein in the *Saccharomyces cerevisiae* ubiquitin-proteasome system. By enabling practical proteome-scale interaction screening, HInt provides a general framework for uncovering hidden homologues and expands the conceptual landscape of protein homology inference.

## MAIN

Despite decades of genomic research, a substantial fraction of proteins across all kingdoms of life remains poorly annotated or entirely uncharacterised. Classical homology detection relies primarily on sequence similarity (BlastP)^1^ and, more recently, structural similarity (Foldseek)^2^. However, evolutionary divergence progressively erodes these signals, often beyond detectable thresholds, while functional relationships between interacting proteins can be maintained. As a result, current approaches lose sensitivity with increasing evolutionary distance, leaving a substantial fraction of homologous relationships unresolved.

Protein-protein interactions (PPIs) provide a complementary and largely untapped source of evolutionary information. Interaction interfaces can remain conserved even when global sequence and structural similarity are no longer detectable^3,4,5,6^. This suggests that interaction interfaces may encode a more persistent evolutionary signal, offering an alternative route to infer functional relationships among highly divergent proteins. However, experimental mapping of PPIs at proteome scale remains impractical, and thus this dimension of homology has remained largely inaccessible.

Recent advances in deep learning-based structure prediction, including AlphaFold^7^ and RoseTTAFold^8^, now enable accurate *in silico* modeling of protein complexes, providing a potential route to systematically explore interaction space. Yet, their application at the scale of entire proteomes remains computationally prohibitive. Existing pipelines, such as AlphaPulldown^9^ and PPIFold^10^, rely on sequential or loosely parallelised execution strategies that limit throughput and restrict genome-wide exploration of interaction networks. In parallel, lightweight PPI prediction methods have been developed to reduce computational cost^11,12,13^, although they generally do not match the accuracy of structure-based approaches across diverse biological systems. In addition to AlphaPulldown^9^ and PPIFold^10^, other AlphaFold-based frameworks, including AlphaFast^14^ and AF_cache^15^, improve throughput by reusing intermediate files or optimising MSA generation through caching and batching strategies. However, these approaches provide limited acceleration of the structure prediction stage itself and do not resolve the computational bottleneck associated with genome-scale PPI scoring.

Here we asked whether conserved interaction interfaces could provide an independent signal for homology detection beyond sequence and structure. To address this question, we developed HInt (Homology by Interaction) (Fig. 1a), an accelerated AlphaFold-based framework that enables practical proteome-scale PPI prediction and allows interaction-based similarity to be explored as a third axis of homology detection. HInt enables scalable, proteome-wide inference of protein-protein interactions by integrating biologically informed pre-filtering with optimised, high-throughput structure-based modelling. Through accelerated MSA generation, adaptive multi-GPU scheduling with memory-aware batching, and robust interaction scoring, HInt enables practical proteome-scale protein-protein interaction screening.

**Fig. 1.**
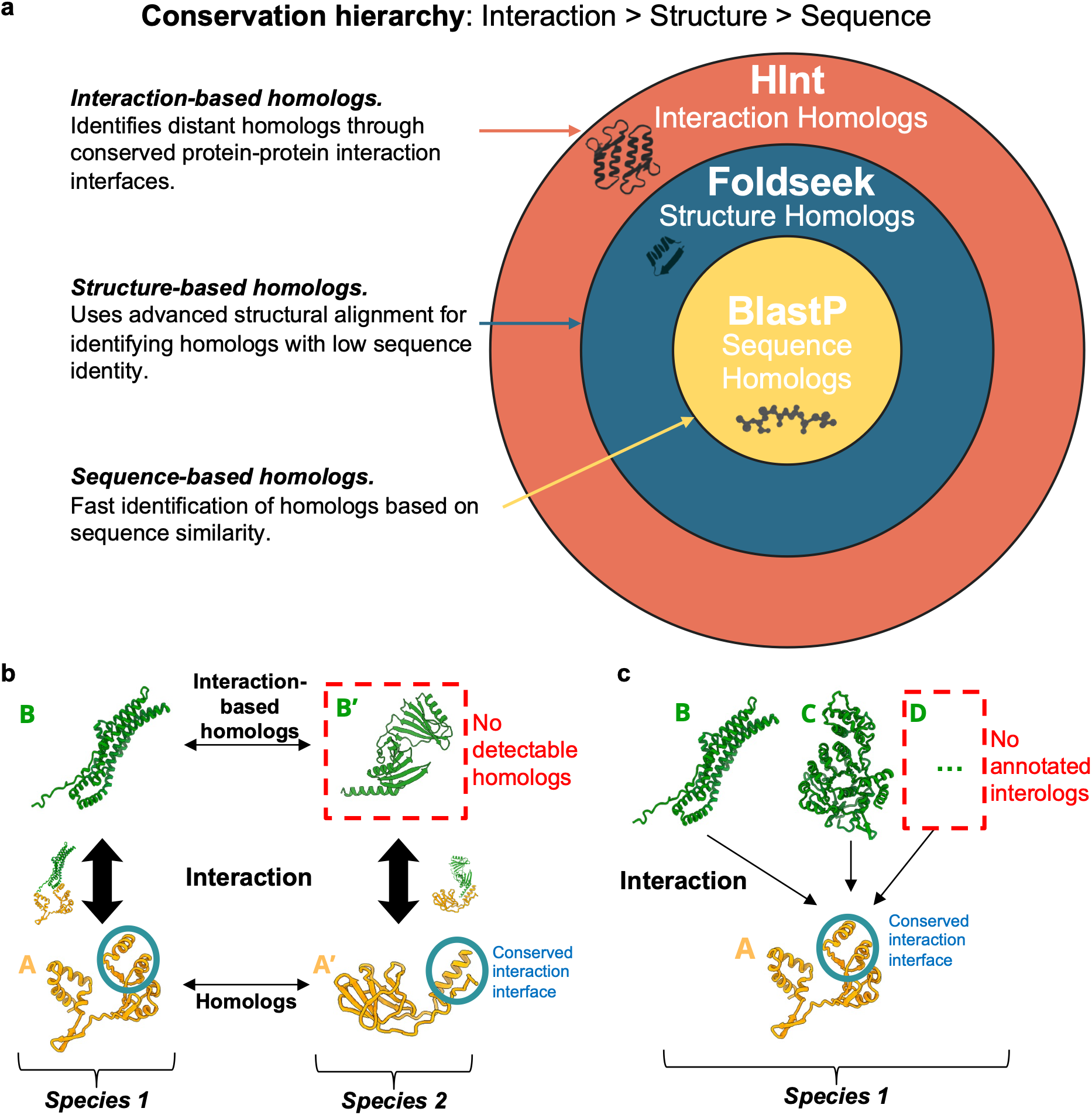
| Interaction-based homology expands the detectable homolog space. **a, Conceptual representation of the hierarchy of homolog detection methods.** The figure illustrates the fraction of homologs identified by different approaches. Sequence- and structure-based methods such as BlastP and Foldseek capture a large portion of the detectable homolog space, whereas interaction-based homology enables the identification of additional homologs with divergent sequences and structures, expanding the overall coverage. **b, Schematic representation of interaction-based homology.** The figure illustrates the relationship between proteins B and B′, defined through their interactions with homologous proteins A and A′. The conservation of interaction partners across homologous proteins supports the inference of interaction-based homology. **c, Schematic representation of an interolog group.** The figure shows an interolog group composed of proteins B and C, defined by their shared interaction with protein A. The conserved interaction interface on protein A is highlighted (blue circle), indicating a shared binding site across the interaction group. Protein D represents an interolog not yet annotated.

Using this framework, we distinguish two types of interaction-based homology: (i) *remote homologues*, in which proteins lacking detectable sequence or structural similarity are identified through a conserved interaction interface with their respective homologous partners (Fig. 1b), and (ii) *interologs*, in which distinct proteins within the same organism interact with a common partner through the same interfaces (Fig. 1c).

We demonstrate the power of this approach through applications to both prokaryotic and eukaryotic systems, supported by experimental validation. In bacteria, HInt identified a previously undetected VirB5 homologue in the F-plasmid type IV secretion system, despite the absence of detectable sequence or structural similarity. In eukaryotes, it revealed a previously unannotated F-box like protein within the ubiquitin-proteasome system. Together, these results establish interaction-based similarity as a robust and exploitable signal for homology detection, uncovering evolutionary relationships that are invisible to conventional approaches.

## RESULTS

### High-throughput pipeline for large-scale AlphaFold-based PPI screening

#### Pipeline overview

HInt enables interaction-based similarity inference through scalable, proteome-wide structure prediction. It is a modular computational framework integrating biological prior knowledge and optimised computational strategies to enable large-scale PPI analysis. The pipeline accepts a bait sequence, optional biological annotations, and a proteome-wide set of candidate proteins. It then performs localisation prediction, structure modeling, and interaction scoring in a fully automated and checkpointed workflow (Fig. 2a). Combined with GPU-aware scheduling and memory optimisation, HInt efficiently processes large protein datasets and produces a ranked list of interaction candidates.

**Fig. 2.**
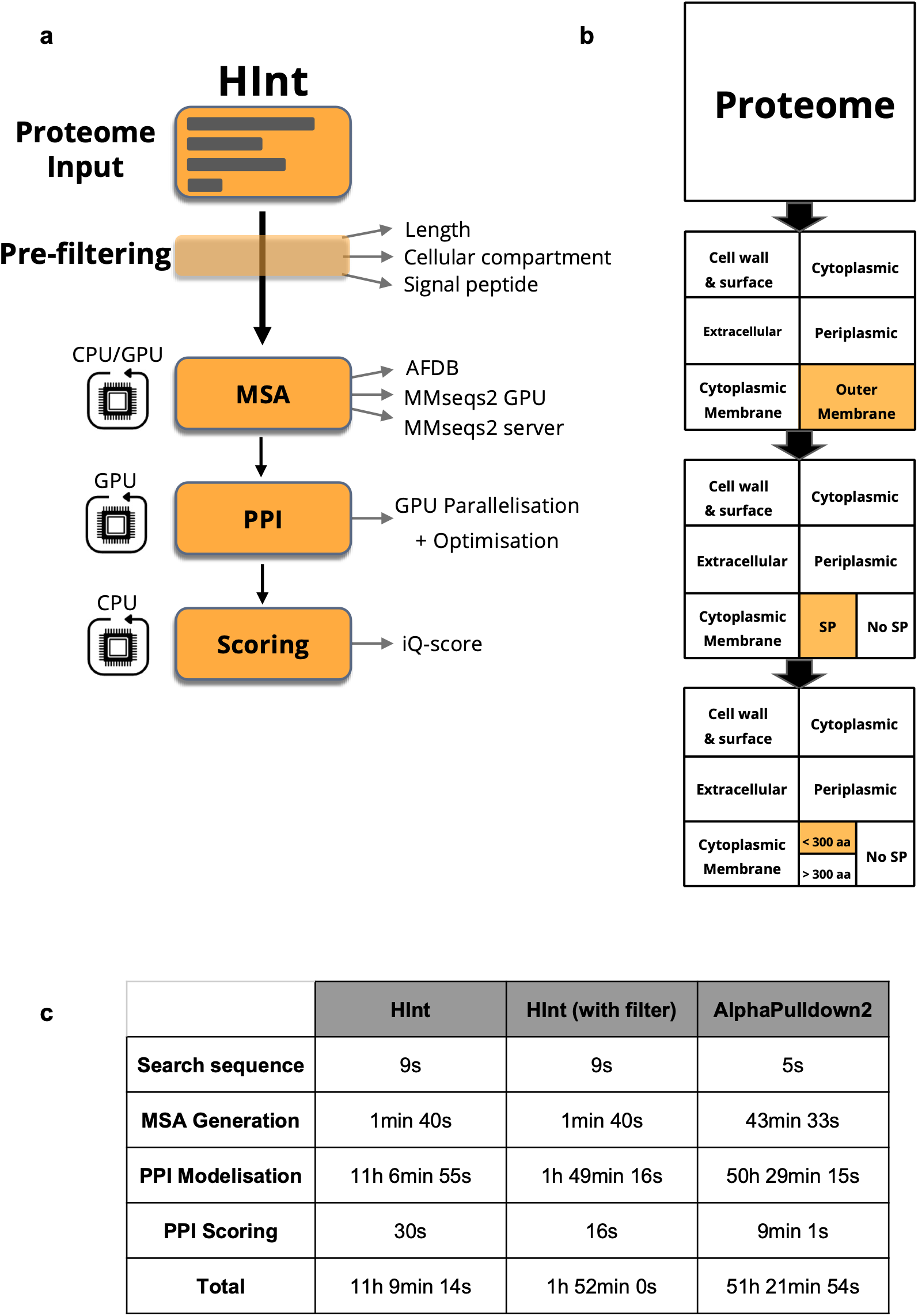
| HInt enables scalable proteome-wide prediction of protein–protein interactions. **a, Overview of the HInt pipeline.** HInt comprises four sequential stages: (i) biologically informed pre-filtering to reduce the interaction search space, (ii) multiple sequence alignment (MSA) generation and feature preparation, (iii) structure-based protein–protein interaction prediction using state-of-the-art deep learning frameworks, and (iv) interaction quality scoring using the iQ-score. GPU- and CPU-aware optimisations are integrated throughout the workflow to enable scalable proteome-wide screening. **b, Biologically informed pre-filtering strategy.** To reduce the combinatorial interaction space before structural prediction, HInt sequentially filters candidate protein pairs according to three biological criteria: predicted cellular localisation, the presence of a signal peptide, and protein length. The example shown selects outer membrane proteins containing a signal peptide and shorter than 300 amino acids. **c, Computational performance comparison.** Execution times are compared for HInt and the AlphaPulldown2 Snakemake workflow using the same benchmark dataset, consisting of 108 PPIs generated by screening P33790 against the complete F-plasmid proteome. HInt is shown both with and without the biologically informed pre-filtering strategy. The average combined sequence length of the protein pairs was 1,135 amino acids.

#### Biologically informed pre-filtering reduces the search space

Genome-wide PPI identification is limited by the combinatorial explosion of one-versus-all screening, which HInt mitigates through a dedicated pre-filtering stage (Fig. 2b). This stage applies sequential biologically informed filters based on protein length, signal peptide detection (SignalP5^16^), and subcellular localization prediction (DeepLoc2^17^/DeepLoc Pro^18^), progressively reducing the candidate search space prior to MSA generation and structure prediction (Supplementary Information 1.1-1.3). All filters are user-configurable to balance sensitivity and computational efficiency, and retained proteins are automatically passed to downstream modeling steps.

#### Accelerated MSA generation

Accurate MSAs are a critical prerequisite for reliable structure-based modeling. Despite prior filtering, the number of candidate proteins can remain substantial, making this step a major bottleneck. HInt addresses this limitation through two complementary strategies (Supplementary Fig. 1a). First, when UniProt identifiers are available, precomputed alignments are retrieved directly from the AlphaFold Database^19^, enabling efficient reuse of existing multiple sequence alignments. Second, for proteins lacking precomputed MSAs, HInt employs GPU-accelerated MMseqs2^20^. Together, these strategies substantially reduce computation time, achieving an average 47-fold speedup compared with the standard MMseqs2 ColabFold MSA workflow (server-based)^21,22^ and an 11-fold speedup over standalone GPU-accelerated MMseqs2 searches (Supplementary Fig. 1b), reflecting both hardware acceleration and reduced query complexity ( Supplementary Information 2.1-2.3). HInt also monitors alignment depth and flags proteins with shallow MSAs (<100 sequences), where limited sequence diversity may compromise prediction confidence. A single MSA is generated per protein and reused across runs *via* local caching, reducing redundant computations and improving efficiency (Supplementary Information 2.4).

#### Dynamic GPU scheduling for genome-scale PPI prediction

Structure-based PPI prediction using AlphaFold-Multimer^23^ and AlphaFold 3^24^ achieves strong performance but remains limited by high computational cost^25^. In standard implementations, predictions are executed sequentially, severely constraining throughput for large-scale interaction screens. HInt addresses this limitation through two complementary optimisation strategies combined with dynamic scheduling (Supplementary Fig. 2a). First, complex structure prediction tasks are distributed across all available GPUs. Second, HInt implements VRAM-aware batching by estimating memory requirements as a function of complex size, thereby maximizing utilisation of each GPU (Supplementary Fig. 2b). This intra-GPU optimisation is particularly effective when using high-memory hardware or modeling large numbers of small complexes. A dynamic job queue continuously monitors and reallocates resources as predictions complete, ensuring near-maximal proper hardware utilisation (Supplementary Information 3.1-3.3). The combination of inter-GPU parallelisation and intra-GPU memory optimisation results in an average 4.5-fold reduction in modeling time (Supplementary Fig. 2c) for AF-Multimer (v2.3.2) and AF3 (v3.0.1), enabling genome-scale PPI screening that would otherwise be computationally prohibitive (Supplementary Information 3.4-3.5). Importantly, these computational gains are achieved without compromising prediction quality.

#### Interaction scoring

Accurate scoring of predicted PPIs is essential for reliable large-scale screening. To identify the most discriminative metric, we systematically benchmarked a range of available approaches, including recent state-of-the-art methods such as pi-score^26^, pDockQ^27^, pDockQ2^28^, iQ-score^10^, ipSAE^29^, LIS^30^, iLIS^31^ and actifpTM^32^. The benchmark was performed on a balanced dataset of 1,000 PPIs comprising 500 experimentally validated positive interactions from IntAct and 500 negative interactions from Negatome. All protein complexes were predicted using AlphaFold-Multimer, and every scoring metric was evaluated on the same set of predicted structures to ensure a fair comparison (Supplementary Information 5.1-5.2). Among the evaluated methods, iQ-score showed the highest discriminative performance (Supplementary Fig. 3a) and was therefore selected as the default scoring metric. Finally, to support high-throughput analyses, HInt parallelises PPI scoring across multiple CPU cores (Supplementary Information 5.3).

#### Pipeline output

The primary output of HInt is a ranked summary table of predicted PPIs based on the selected scoring metric. For each interaction, the table reports the iQ-score, along with relevant annotations including predicted subcellular localisation and presence or not of signal peptide. To facilitate interpretation, each prediction is accompanied by representations, including distograms, amino acid contact matrices, and the AlphaFold model of the complex, with interface residues automatically highlighted based on their predicted contribution to the interaction. HInt additionally generates a summary table reporting the filtering status of each candidate protein, together with a timing report detailing the execution time of each pipeline step (Supplementary Information 6). Together this allows biologists to easily analyse HInt results and select candidate protein and amino-acids for mutation and experimental validation.

#### End-to-end pipeline performance evaluation

To evaluate the performance of the complete HInt pipeline, we measured the execution time required for MSA generation, structure prediction and interaction scoring, and compared it with the state-of-the-art AlphaPulldown2 pipeline (Snakemake implementation v2.1.8). No pre-filtering was applied, requiring both pipelines to generate all MSAs, build all structural models and score all predicted interactions. The comparison was performed using MMseqs2 for AlphaPulldown2, 64 CPU cores and four NVIDIA RTX 4500 Ada GPUs for both pipelines (Supplementary Information 7). Under these conditions, HInt completed the analysis five times faster than AlphaPulldown2 while producing the same set of predicted PPIs. When combined with HInt’s biologically informed pre-filtering strategy, the overall runtime was further reduced, resulting in a 25-fold acceleration (Fig. 2a) (Supplementary Information 8.1 for filters).

### Experimental validation of interaction-based homology

To validate the conceptual framework underlying HInt, we selected one representative example for each type of interaction-based homology. We first investigated remote homologue discovery in bacterial type IV secretion systems, followed by interolog discovery in the eukaryotic SCF ubiquitin ligase complex.

#### Experimental validation of divergent homologues identification

We first exploited the bacterial type IV secretion systems (T4SSs), versatile nanomachines that exhibit exceptional diversity^33^. Encoded by mobile genetic elements and widespread across bacteria, T4SSs display extensive sequence and structural divergence. Despite this variability, they share a conserved set of 12 core components, designated VirB1-VirB11 and VirD4, which are essential for function. Notably, extensive sequence- and structure-based searches, the reference and well-characterised F-plasmid system lacks one of these core proteins, VirB5^32^, illustrating the limits of current homology detection approaches.

The VirB5 protein forms the tip^34^ of the filament and specifically interacts with its VirB6 partner in the stalk complex; both proteins assemble into homopentameric complexes^35^ (Fig. 3a). Building on these properties, we applied HInt to all proteins encoded by the F-plasmid, using TraG_B6_ as bait and incorporating prior knowledge of VirB5 features, namely periplasmic/extracellular localisation and the presence of a signal peptide.

**Fig 3.**
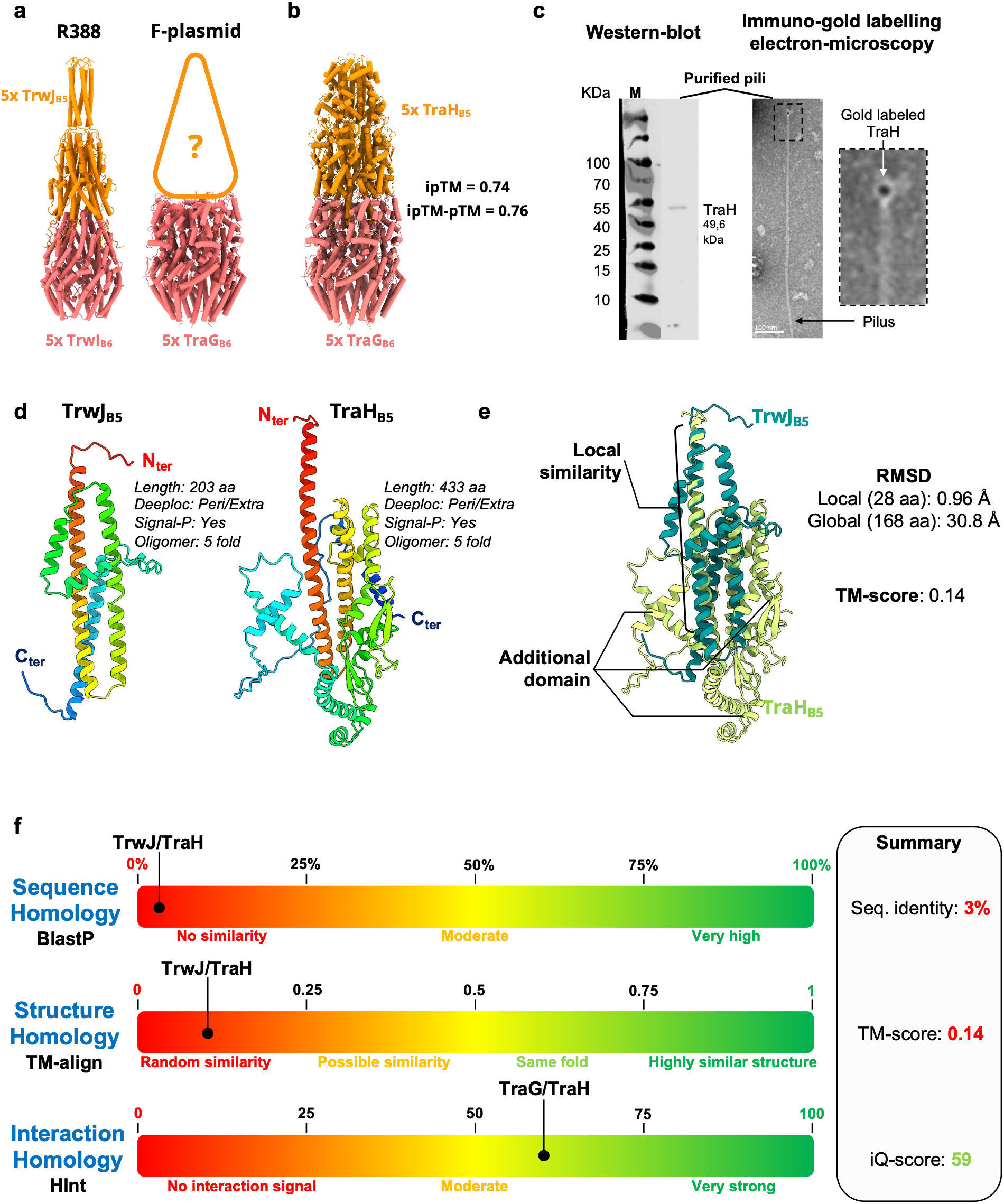
| Interaction-based homology identifies TrwJ_B5_ and TraH_B5_ as remote homologs despite undetectable sequence and structural similarity. **a, Structural context of the conjugative Type IV secretion system (T4SS) stalk complex.** Left, AlphaFold 3 model of R388 stalk complex based on PDB 8RT9 containing five copies of TrwJ_B5_ (orange) and the pentameric VirB6 homolog TrwI_B6_ (red). Right, AlphaFold 3 model of the F-plasmid VirB6 homolog TraG_B6_ assembled as a pentamer (red). The orange outline above the complex represents the unknown F-plasmid tip protein whose identity is inferred through interaction-based homology. The intrinsically disordered C-terminal region of TraG_B6_ (residues 452-938) was not modeled. **b, AlphaFold 3 model of the predicted F-plasmid stalk complex.** Predicted stalk complex composed of pentameric TraG_B6_ (red) and five copies of TraH_B5_ (orange). The intrinsically disordered C-terminal region of TraG_B6_ (residues 452-938) was omitted from the model. **c, Experimental validation of TraH_B5_ localisation at the F-pilus tip.** Western blot analysis of purified pili and immunogold electron microscopy confirm the presence of TraH at the distal pilus tip. **d, AlphaFold 3 models of TrwJ_B5_ and TraH_B5_ monomers**. Although the two proteins share a similar arrangement of α-helices, their overall architectures differ substantially, notably due to the presence of an additional domain in TraH_B5_. **e, Structural superposition of TrwJ_B5_ and TraH_B5_.** Global structural alignment reveals no significant similarity (TM-score = 0.14), although a short α-helical region can be locally superimposed, suggesting a shared ancestral structural element between TrwJ_B5_ and TraH_B5_. **f, Comparison of sequence-, structure-, and interaction-based similarity metrics.** TrwJ_B5_ and TraH_B5_ share only 3% sequence identity and display no detectable global structural similarity (TM-score = 0.14). In contrast, HInt assigns a strong interaction similarity score (iQ-score = 59), revealing a remote homologous relationship that is undetectable using conventional sequence- or structure-based approaches. Thus, only interaction-based homology identifies TrwJ_B5_ and TraH_B5_ as remote homologs.

From 110 proteins, HInt identified TraH (Uniprot ID P15069) within 1 hour 52 minutes (Fig. 2c) as a candidate for the F-plasmid VirB5 homologue (Fig. 3b, Supplementary Fig. 4a, Supplementary Information 8.1). Given that VirB5 forms the filament tip^34^, we experimentally tested this prediction by assessing the localisation of TraH. To this end, we introduced a streptavidin tag into TraH, isolated F-T4SS filaments, and performed both western blot and immunogold labelling. As shown in Fig. 3c, TraH is detected in purified filament samples by western blot and is specifically localised at the filament tip by negative-stain electron microscopy using immunogold labelling.

Strikingly, TraH_B5_ shares less than 3% sequence identity with previously characterised VirB5 proteins. Although the N-terminal helix-loop-helix (HLH) motif is locally conserved, structural comparisons at the monomer level reveal no meaningful global similarity (RMSD >30 Å), indicating that this local conservation is insufficient for homology detection (Fig. 3d,e). In contrast, when considered as part of their respective complexes, the predicted pentameric TraH_B5_-TraG_B6_ assembly recapitulates the overall architecture of the canonical TrwJ_B5_-TrwI_B6_ complex (Fig. 3a). These findings indicate that HInt captures conserved interaction-level organisation, enabling the identification of functional homologues that remain undetectable by sequence- or structure-based approaches alone (Fig. 3f).

#### Experimental validation of interolog identification

In eukaryotes, targeted protein degradation is largely mediated by the ubiquitin-proteasome system, which relies on ubiquitin ligases (E3s) to selectively mark substrates for degradation. A central example of E3 is the SCF (Skp1/Cullin-1/F-box) complex. It is a modular assembly in which the Skp1/Cullin-1 core acts as a structural scaffold, while F-box proteins function as interchangeable substrate receptors and confer specificity to the complex (Fig. 4a,b). This organisation enables a single core complex to recognise a wide diversity of targets through a repertoire of F-box proteins, each containing a conserved F-box domain that mediates binding to Skp1.

**Fig. 4.**
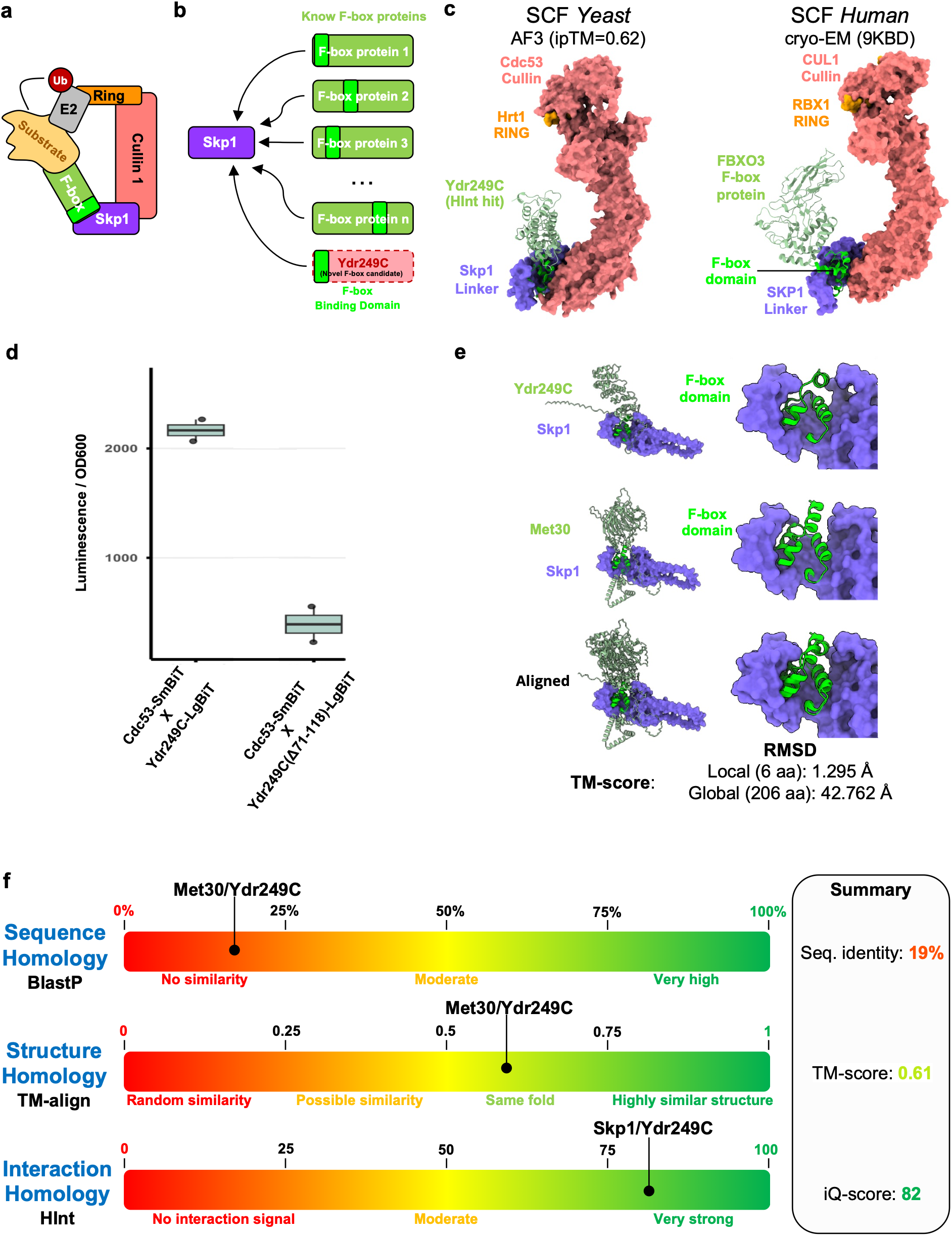
| Discovery and validation of a divergent F-box protein using interaction-based homology. **a, Organisation of the SCF (Skp1/Cullin/F-box) ubiquitin ligase complex.** The Cullin-RING module recruits the ubiquitin-charged E2 enzyme, whereas F-box proteins act as substrate adaptors through a conserved F-box domain that mediates binding to Skp1. **b, Diversity of F-box proteins.** Schematic illustrating the diversity of F-box proteins, whose conserved F-box domain can be embedded within highly divergent sequence contexts and domain architectures, limiting its detection by conventional sequence-based approaches. **c, Structural comparison between the SCF complex predicted by HInt and the cryo-EM structure of the human Cullin–Rbx complex.** AlphaFold 3 model of the yeast SCF complex incorporating Ydr249C (left), compared with the experimentally determined human SCF complex (PDB 9KBD; right). The predicted complex recapitulates the canonical SCF architecture and positions the N-terminal region of Ydr249C at the Skp1 interface, consistent with an F-box-mediated interaction. **d, NanoBiT validation of Ydr249C incorporation into an SCF complex.** Ydr249C generates a robust NanoBiT signal with Cdc53, whereas deletion of the predicted F-box-like region or mutation of Skp1 markedly reduces the signal, demonstrating that this region is required for incorporation into an SCF complex. **e, Structural comparison of the Skp1 complexes formed by Ydr249C and the canonical F-box protein Met30.** Close-up views of the interaction interfaces reveal that Ydr249C adopts an α-helical motif and interaction geometry similar to those of Met30. Based on structural inspection, the F-box-like region spans residues 71-118 in Ydr249C and residues 183-231 in Met30. **f, Comparison of sequence-, structure-, and interaction-based similarity metrics.** When restricted to the F-box-like region, Ydr249C shares only 19% sequence identity with the canonical F-box protein Met30 and exhibits only moderate local structural similarity (TM-score = 0.61). In contrast, HInt assigns a strong interaction score (iQ-score = 82), correctly identifying Ydr249C as a divergent F-box protein.

F-box domains are short (~40-50 amino acids), predominantly α-helical motifs that mediate interaction with Skp1, annotated in InterPro by IPR001810 (F-box domain) and IPR036047 (F-box-like domain superfamily) entries^36^. Despite their conserved function, these domains are degenerate in sequence and enriched in common hydrophobic residues, resulting in low information content^37,38^. This limitation is further amplified by the embedding of the F-box domain within variable protein architectures, where divergence in sequence context and domain organisation can obscure its detection. As a result, the two InterPro entries are not sufficient to annotate the full repertoire of F-box proteins^36^ and it is possible that further F-box like substrate receptors remain to be identified.

We used HInt to screen the entire *S. cerevisiae* (GenBank assembly GCF_000146045.2) cytoplasmic and nucleus proteome against Skp1. Within 8 days, HInt analysed 6033 candidate proteins and produced a ranked list of potential F-box-containing proteins (Supplementary Fig. 5a, Supplementary Information 9.1). Strikingly, 20 of the 22 top-ranking hits are known F-box proteins^39^, including 6 that are not annotated with IPR001810 nor IPR036047, thus confirming the high sensitivity and specificity of the approach. Notably, the highest-ranking unannotated protein appeared at rank 2 (Ydr249C).

The yeast protein Ydr249C (Uniprot ID Q03787) shows no detectable match to the canonical F-box domain (IPR001810, IPR036047). However, the AlphaFold model of the Skp1/Ydr249C complex revealed a binding mode consistent with canonical Skp1-F-box interactions. When modelled in the context of the full SCF complex, Ydr249C adopts a configuration compatible with known SCF architectures (Fig. 4c), suggesting that it can integrate into the complex in a manner similar to established F-box proteins.

We next performed experimental validation to determine whether Ydr249C can be incorporated into SCF complexes. Using a NanoBiT complementation assay, which enables sensitive detection of protein-protein interactions of endogenously expressed proteins in *S. cerevisiae*^41^, we tested the interaction between the yeast Cullin-1 (Cdc53) and Ydr249C (Fig. 4d, supplementary Fig. 5b). The NanoBiT assay revealed a clear interaction signal between Cdc53 and Ydr249C, which is abrogated by deleting the F-box-like domain of Ydr249C (residues 71-118) or in a *skp1-3* mutant. Together, these results demonstrate that the previously uncharacterised protein Ydr249C is an F-box protein, discovered by HInt and experimentally validated.

To further compare this newly identified F-box protein with canonical members of the family, we analysed its predicted F-box region alongside that of Met30, whose interaction with Skp1 has been well described^40^ (Fig. 4e). Pairwise alignment of the candidate region revealed poor sequence identity (~19%) over ~47 residues. Structural comparison showed a similar fold, with a TM-score of 0.61, indicating that both segments adopt a similar α-helical architecture characteristic of F-box domains. However, despite this local similarity, the region is not detected by structural-based similarity methods (Fig. 4f).

Together, these results establish that the previously uncharacterised protein Ydr249C is an F-box protein identified by HInt and experimentally validated. We therefore propose naming this newly identified protein Fbl1 (F-box-like protein 1). More broadly, these findings demonstrate that interaction-based similarity can uncover homologous proteins that escape conventional sequence-and structure-based annotation. In protein families where sequence conservation has largely eroded, conserved interaction interfaces provide a powerful evolutionary signal for homology detection.

## DISCUSSION

Our findings establish interaction-based similarity as a third axis of homology detection, complementing sequence- and structure-based approaches. By demonstrating that conserved interaction interfaces can reveal evolutionary relationships beyond the reach of conventional homology detection, this work expands the conceptual framework used to infer protein homology.

To test this hypothesis, we developed HInt, an accelerated AlphaFold-based framework enabling practical proteome-scale interaction screening. Through biologically informed filtering, optimised MSA generation and efficient multimer prediction, HInt transforms genome-wide PPI prediction from a computationally prohibitive task into a practical discovery strategy. These optimisations are broadly applicable to deep learning-based structural biology pipelines and are expected to benefit further from continuing advances in AI models and GPU hardware.

Validation in two unrelated biological systems illustrates the discovery power of interaction-based similarity. HInt identified a previously unknown VirB5 homologue (TraH_B5_) in the F-plasmid type IV secretion system and uncovered Ydr249C as a previously unrecognised F-box protein, both of which were subsequently validated experimentally. These findings indicate that interaction interfaces may remain under evolutionary constraint even after sequence and global structural similarity have become undetectable, allowing conserved molecular functions to be recovered through interaction interfaces alone.

Rather than replacing existing approaches, interaction-based similarity complements sequence- and structure-based homology detection by providing access to a distinct layer of evolutionary information. Sequence conservation remains highly effective for closely related proteins, structural similarity extends homology detection across greater evolutionary distances, whereas interaction-based similarity becomes particularly informative once both signals have largely eroded. Because HInt relies on accurate multimer prediction, its performance will continue to improve alongside advances in AI-based interaction modelling.

Interaction-based similarity is not intended to replace established sequence- and structure-based approaches, which remain considerably faster, highly robust and benefit from decades of methodological development and benchmarking. Instead, HInt is designed for situations in which these approaches reach their limits. Although interaction-based modelling remains computationally demanding and depends on the accuracy of predicted protein complexes, it provides access to a distinct evolutionary signal that becomes particularly informative in the “twilight zone” of homology, where sequence and structural similarity are no longer detectable but conserved interaction interfaces persist. At present, such analyses are likely to remain targeted towards specific biological questions or challenging protein families rather than routine genome annotation. However, continuing improvements in AI-based structure prediction and hardware acceleration are expected to progressively reduce these computational barriers, making interaction-based homology detection increasingly accessible.

Beyond enabling the discovery of individual proteins, our results suggest that conserved interaction interfaces represent a complementary organising principle of protein space that can refine genome annotation, improve protein family classification and reveal evolutionary relationships hidden from conventional approaches. More broadly, our findings support the view that protein interactions constitute an evolutionary signal in their own right, establishing interaction-based similarity as a complementary axis for exploring the most divergent regions of protein evolution.

## SUPPLEMENTARY INFORMATION

Methods and ED Figures 1-5.

## CODE AVAILABILITY

The HInt source code and installation instructions are available at : https://github.com/Qrouger/HInt.

## ACKNOWLEDGEMENTS

We thank Y. Lefrancois, M. Choukair, O. Delalande for software test. EM data were collected at TEM2C (Biosit UAR 3480 US18).

## ETHIC DECLARATION

The authors declare no competing interests.

## CONTRIBUTIONS

Q.R., E.G., D.F.M. and K.M. conceived the study and contributed to the development of the conceptual framework. Q.R. developed and implemented HInt. P.P. and M.T. performed the biological experiments for the identification and validation of the VirB5 homolog. G.R. conceived the experiments related to the identification of new F-box proteins, which were performed and analysed by E.T. D.F.M and K.M supervised the project and all authors contributed to writing and revising the manuscript.

