## Supplementary Information for "HInt: interaction-based homology discovery through accelerated genome-scale AlphaFold screening"

#### METHODS

#### 1. Installation, Input parameters and protein pre-filtering

HInt is designed for genome-wide one-versus-all protein–protein interaction (PPI) modeling. The software is distributed as a Conda package, enabling straightforward installation through the provided installation script. To make proteome-scale screening computationally feasible, HInt incorporates a series of biologically informed pre-filtering steps that restrict the candidate interaction space before computationally intensive alignment and structure prediction. Protein sequences are retrieved either from an input FASTA file or from UniProt identifiers and are stored locally in a serialized dictionary to enable efficient access and reuse throughout the pipeline.

##### 1.1 Protein length filtering

The first filter applied is protein length. By default, proteins shorter than 20 amino acids are excluded to remove short peptide fragments that are unlikely to fold into stable protein domains.

Users can define both the minimum and maximum protein lengths according to their screening strategy. Proteins falling outside these user-defined thresholds are removed from the candidate set prior to downstream analysis. This initial filtering step eliminates a substantial fraction of annotated open reading frames, thereby reducing the computational burden while retaining biologically relevant candidates.

##### 1.2 Subcellular localisation prediction

Depending on the organism, HInt performs subcellular localisation prediction using DeepLoc2 (v2.0)[17] for eukaryotic proteomes or DeepLoc Pro (v1.0.0)[18] for prokaryotic proteomes. This step is executed systematically, even when no localisation filter is explicitly specified, ensuring that all proteins are annotated with predicted cellular compartments in the output results. Localisation predictions are cached locally to enable reuse across independent runs. When a specific cellular compartment is defined as a constraint (*e.g.*, extracellular, membrane, cytoplasmic…), HInt restricts the candidate list to proteins predicted to localise to the specified compartment. Filtering constraints eliminate biologically implausible interaction pairs.

##### 1.3 Signal peptide prediction and sequence trimming

Signal peptide prediction is performed using SignalP5 (v5.0b)[16] for all candidate proteins. Predicted signal peptides serve a dual purpose within HInt. First, when present, the signal sequence is removed to retain only the mature, biologically relevant region for downstream modeling. This trimming step additionally reduces effective protein length, which directly impacts GPU memory requirements during structural modelling. Second, signal peptide annotations are cached locally and can be used as a filtering criterion: when signal peptide status is specified as a constraint, only proteins matching the defined condition are retained in the candidate prey set. This is particularly relevant when screening for secreted or surface-exposed interaction partners in bacterial systems.

##### 1.4 Bait protein configuration

A bait protein can be specified using the Interact_with parameter. This protein defines the interaction target against which all candidate proteins are screened. HInt supports three complementary bait configurations that provide increasing specificity:

**Interface-restricted screening.** Users may optionally restrict the interaction search to specific regions of the bait protein by defining residue intervals corresponding to the expected interaction interface. This enables faster and more precise interaction screening.

**Sequential multi-bait screening.** Multiple bait proteins can be specified. When several bait proteins are provided, all candidate prey proteins are first evaluated for interaction with the primary bait. Only prey proteins predicted to form a complex with a low inter-PAE with this bait are carried forward and evaluated against the second bait, and so on. Consequently, the order of the bait proteins determines the sequence of the filtering steps and therefore their relative priority.

**Multimeric bait.** Alternatively, a multimeric bait can be specified by providing multiple proteins known to interact together. In this configuration, the proteins are modeled as a single complex and used collectively as a unified bait during interaction screening.

##### 1.5 Configurable filter stringency

Because overly strict constraints could exclude true interactions, HInt is designed to allow flexible configuration of these filters, ensuring a controlled balance between computational efficiency and sensitivity.

#### 2. Multiple sequence alignment generation

##### 2.1 Benchmarking MSA generation strategies

Multiple sequence alignment (MSA) generation is a critical component of the pipeline, directly impacting computational efficiency and downstream structural modeling quality. To reduce runtime while preserving alignment depth, HInt integrates complementary state-of-the-art strategies depending on input availability and dataset size (Supplementary Fig. 1a).

To compare the runtime of different MSA generation strategies, we constructed six biologically relevant protein datasets with varying numbers of sequences to reflect realistic screening conditions (Extended Data). Each dataset was processed using 48 CPU and 1 GPU RTX 6000 ada. The computational time required for each method was recorded (Supplementary Fig. 1b).

##### 2.2 Precomputed MSA retrieval from AFDB

When UniProt identifiers are provided, HInt retrieves precomputed MSAs (A3M format) from the AlphaFold Database (AFDB)[19]. These alignments correspond to the full-length protein sequence. If a signal peptide is predicted, the sequence is trimmed to retain only the mature region, and the MSA is realigned using MAFFT (v7.490)[42] to ensure positional consistency with the processed sequence. Previously generated MSAs are automatically detected and reused when available, reducing computational cost and accelerating subsequent analyses.

##### 2.3 *De novo* MSA generation using MMseqs2

If UniProt identifiers are not provided, and if more than ten proteins lack precomputed MSAs, HInt performs de novo alignment generation using GPU-accelerated MMseqs2 (v1.6.1)[20]. For small numbers of remaining sequences (≤10 proteins), the MMseqs2 ColabFold server (v1.5.5)[21,22] implementation is used instead. In both cases, MSAs are generated from the processed mature sequences (i.e., signal peptides removed when present).

##### 2.4 MSA saving and quality monitoring

All generated MSAs are stored locally and reused in subsequent screening runs to avoid redundant computation. Alignment quality is monitored through the computation and visualisation of MSA depth metrics. For each protein, the number of aligned sequences is reported. Proteins associated with shallow MSAs (fewer than 100 sequences) are automatically flagged and recorded for user inspection. Limited evolutionary depth may reduce the confidence and interpretability of downstream PPI predictions.

### 3. PPI prediction: GPU optimisation and VRAM-aware scheduling

HInt accelerates large-scale protein–protein interaction (PPI) prediction by combining two complementary optimisation strategies with a dynamic GPU scheduling system. For each PPI, five models are generated, with up to three recycling iterations per model.

##### 3.1 Inter-GPU parallelisation

The first optimisation relies on parallelisation across multiple GPUs. PPI modeling jobs are distributed across all available GPUs, allowing multiple interactions to be predicted simultaneously. Each GPU processes independent PPI prediction tasks, enabling acceleration when additional GPUs are available. For example, when four GPUs are used, up to four independent PPI predictions can be executed in parallel (Supplementary Fig. 2a).

##### 3.2 Intra-GPU optimisation

The second optimisation focuses on GPU memory (VRAM) utilisation. The amount of VRAM reserved by AlphaFold during inference depends primarily on the combined length of the interacting proteins. We benchmarked this relationship by generating different length PPI on RTX 6000 ada (48 Gb) and RTX 4500 ada (24 Gb) using AlphaFold-Multimer (v2.3.2)[23]. JAX execution was configured using the following settings to optimise GPU memory management: XLA_PYTHON_CLIENT_PREALLOCATE=false,TF_FORCE_UNIFIED_MEMORY=true , and xla_gpu_enable_triton_gemm=false.

We derived an empirical polynomial model linking sequence length (L, in amino acids) to VRAM usage (V, in Gb) (Supplementary Fig. 2b). HInt uses an intentionally conservative estimate of this function to schedule multiple PPI predictions on the same GPU when sufficient memory is available (Supplementary Fig. 2a).

$$V = 3.8 - 6.27 \times 10^{-5} L + 3.32 \times 10^{-6} L^2$$

This approach maximises GPU utilisation while preventing out-of-memory crashes. The impact of this optimisation is particularly significant when using GPUs with large VRAM capacity and when modeling large numbers of small protein interactions.

##### 3.3 Dynamic job scheduling

HInt manages all prediction tasks through a dynamic job queue that continuously optimises GPU workload distribution. Jobs are evenly distributed across available GPUs, and the queue is continuously monitored. When a prediction task finishes, the scheduler immediately assigns the next most suitable job based on the estimated VRAM requirements. This dynamic scheduling strategy enables HInt to fully exploit both parallel GPU execution and optimised memory allocation, thereby maximising overall computational throughput.

##### 3.4 Benchmarking optimisation strategies

To evaluate the impact of the different optimisation strategies implemented in HInt, we compared two configurations: standard execution and HInt execution Using AlphaFold3 (v3.0.1)[24] and AlphaFold-Multimer (v2.3.2)[23] to ensure consistent benchmarking conditions (Supplementary Fig. 2c). Three protein datasets of varying sizes were constructed, and all interaction models were generated using RTX 4500 ada (24Gb) GPU, 64 CPU AMD Ryzen Threadripper PRO 5975WXs. PPI datasets of different sizes were generated such that all interactions present in the smallest dataset were also included in larger datasets, ensuring a nested structure across all benchmark sets.

The dataset of 1,000 PPIs was constructed using protein P33790 from the F-plasmid as bait, paired with randomly selected proteins from the *E. coli* genome (NCBI accession: U00096.3). The datasets of 100 and 10 PPIs were then generated by homogeneous subsampling of the largest dataset, preserving the distribution of PPI lengths across all subsets (~1221,85 aa).

These PPI datasets include interactions of varying sequence lengths, enabling comparison of the different optimisation strategies under conditions that approximate realistic biological scenarios. The resulting modeling times were then compared across configurations.

##### 3.5 Significance of the computational innovations

While structure prediction frameworks such as AlphaFold-Multimer and AlphaFold 3 provide highly accurate structural modeling, they are not designed for large-scale systematic screening of protein–protein interactions. HInt introduces several novel strategies to address this limitation: first, by enabling parallelisation of PPI predictions across multiple GPUs, which allows multiple interactions to be modeled simultaneously; second, by implementing VRAM-aware batching to maximize GPU memory utilisation and prevent out-of-memory crashes; and third, through dynamic job scheduling that continuously optimises GPU workload distribution. Together, these innovations transform single-structure prediction workflows into a scalable, high-throughput platform specifically tailored for genome-wide PPI discovery.

#### 4. Checkpointing and intermediate data persistence

All multiple sequence alignments, figures, scores, datasets, and computationally intensive intermediate results are systematically saved during pipeline execution. HInt implements systematic checkpointing by storing relevant information in a local serialized data structure, enabling robust recovery from interruptions. Upon restart, the pipeline automatically detects existing intermediate results and recomputes only missing or incomplete steps, minimizing redundant CPU and GPU usage. Previously generated data (MSA) can also be reused across runs when analysing different bait proteins within the same dataset, further reducing unnecessary re-computation and enabling efficient iterative screening.

#### 5. PPI scoring and metric benchmarking

Accurate scoring of predicted protein–protein interactions (PPIs) remain a major challenge in the field. Numerous scoring metrics exist, and to identify the most informative measure, we benchmarked all relevant scores using a non-redundant curated dataset of 1,000 PPIs, composed of 500 true positives and 500 true negatives.

##### 5.1 Benchmark dataset construction

The true positive set was derived from experimentally validated interactions in the IntAct database, with high-confidence interactions selected based on IntAct scoring (MI Score >= 0,75). The true negative set was obtained from the Negatome database. All interactions are associated with published references (PMIDs) (Extended Data).

To minimize bias, the dataset includes interactions from multiple species and comprises diverse protein pairs, ensuring that each protein is used only once and excluding homodimeric interactions. Multiple sequence alignments were generated using MMseqs2-GPU (v1.6.1), and structural models were predicted using AlphaFold-Multimer (v2.3.2).

##### 5.2 Metrics benchmarked

For each scoring metric, we applied the thresholds reported in the corresponding publications and evaluated their discriminative performance using standard statistical analyses. Each interaction was evaluated without applying a PAE threshold prior to scoring. Based on these thresholds, predicted interactions were classified as true or false. Sensitivity and specificity were then calculated, and receiver operating characteristic (ROC) curves were generated to assess overall performance (Supplementary Fig. 3a). Area under the curve (AUC) values were computed for quantitative comparison between scoring methods.

##### 5.3 Selection of iQ-score

Based on this benchmark, iQ-score[10] was selected as the default scoring metric for HInt because it achieved the highest overall discriminative performance, as measured by the area under the ROC curve (AUC). Scoring is performed exclusively on the best-ranked model among the five generated for each PPI. An initial filtering step is applied: only models with inter-chain predicted aligned error (PAE) below 10 Å with the prey are considered for scoring, and models exceeding this threshold are excluded from evaluation. When the bait is defined as a multimer, the multimeric assembly is assumed to be known and is therefore not subjected to scoring. Accordingly, the PI-score[26] used in the iQ-score calculation is computed only for interfaces between bait and prey chains. To enable high-throughput scoring, HInt implements CPU-based parallelisation, allowing multiple interactions to be scored simultaneously and substantially reducing overall computation time. Each CPU core independently evaluates the iQ-score of a single predicted protein complex.

#### 6. Output files and visualisation

HInt produces comprehensive output files designed for both systematic analysis and detailed inspection of individual interactions.

**Ranked interaction tables.** All proteins matching the input search list are reported in a file named All_Final_result_HInt.csv, ranked by iQ-score. When multiple bait proteins are specified, the list is ordered based on the cumulative iQ-score across all baits. A complementary file, Summary_result_HInt.csv, provides a summary of the criteria used to include or exclude each protein from the candidate set. HInt_report.txt reports all input parameters used in HInt and the execution time of each pipeline step.

Numerous figures are generated to facilitate in-depth interpretation of predicted interactions.

**Structural visualisation.** For each model passing the PAE cutoff (<= 10Å), a distogram is produced, and a corresponding table summarizes the residues involved in the interaction (PAE and distance <= 10Å), including inter-chain distances and PAE values. Three-dimensional PDB models are annotated by coloring interface residues using the B-factor column. Finally, for each validated protein, a figure depicting interface usage across all validated PPIs is generated, enabling direct comparison of interface regions among different interaction partners.

#### 7. Performance comparison with AlphaPulldown2

To evaluate HInt against the AlphaPulldown2 Snakemake pipeline (v2.1.8), we modeled interactions between protein $TraG_{B6}$ and the complete F-plasmid proteome. Both pipelines were executed using 64 CPU cores and four NVIDIA RTX 4500 Ada GPUs. For AlphaPulldown2, MSAs were generated using MMseqs2 with the --use_mmseqs2 parameter enabled. For both pipelines, models were generated with AlphaFold-Multimer (v2.3.2). A total of 108 PPIs were successfully modeled, excluding two interactions that failed due to out-of-memory (OOM) errors. HInt completed the analysis in 11 h 9 min 14 s, whereas the AlphaPulldown2 Snakemake pipeline required 51 h 21 min 54 s (Fig. 2c). The major differences in computational performance were observed during MSA generation, PPI modeling, and PPI scoring, with HInt achieving a 30-fold acceleration for MSA generation and PPI scoring, and a 5-fold acceleration for PPI modeling, even without biologically informed pre-filtering. When biologically informed pre-filtering was applied, HInt achieved a 25-fold acceleration in PPI modeling by substantially reducing the number of PPI models generated.

#### 8. Experimental validation: Identification of a VirB5 pilus tip protein in the F-plasmid T4SS

The following procedures describe the HInt-based identification and subsequent experimental validation of the divergent VirB5 homologue (P15069) in the F-plasmid type IV secretion system (T4SS).

#### 8.1 Identification of the VirB5 homologue using HInt

HInt was applied to the complete F-plasmid proteome (GenBank accession MK492260.1) using $TraG_{B6}$ (P33790) as the bait protein. Candidate proteins were sequentially filtered based on predicted periplasmic or extracellular localization, reducing the search space from 110 to 46 proteins, and subsequently on the presence of a signal peptide, further reducing the candidate set to 16 proteins. HInt identified TraH (P15069) as a candidate (Supplementary Fig. 4a). Structural visualisation of the predicted interaction interface, combined with the localisation of interface residues, provided additional support for the assignment of TraH as a putative VirB5 homologue.

#### 8.2 Bacterial strains, plasmids and constructs

Mutant strain F-plasmid-Δ*trah* was constructed by lambda Red recombination using pKD3 and pKD46, following the method of Datsenko & Wanner and verified by PCR and sequencing. New constructs were generated using NEBuilder HiFi DNA Assembly kit (New England Biolabs) according to the manufacturer's instructions.

#### 8.3 Purification of F-pilus tip complexes

Native F-pilus tip complexes were purified by batch affinity purification. Briefly, F-pili were sheared from the *F*-plasmid Δ*traH* strain expressing N-terminally Strep-tagged TraH (pBAD-NterSTREP-TraH). The extracellular filament fraction was incubated with Strep-Tactin XT 4Flow resin (IBA Lifesciences) to affinity-purify F-pilus tip complexes through the Strep-tagged TraH protein. Eluted fractions were pooled and concentrated by ultracentrifugation at 300,000 ×

*g* for 2 h. The resulting pellet was resuspended in buffer A (100 mM Tris-HCl, 200 mM NaCl, 10 mM EDTA, pH 8.0).

##### 8.4 Western blotting

Western blotting was performed after protein separation by SDS-PAGE gradient gel 4-20% using standard protocols with STREP-tag antisera (MAN0016358), following the manufacturer protocol and detected with anti-rabbit horseradish peroxidase-conjugated secondary antibody (Bio-Rad), using a chemiluminescence-based detection system (Bio-Rad).

##### 8.5 Negative-stain electron microscopy

Negative stain electron microscopy (NS-EM) was used to analyse the pili-tip complexes. 3 μL of sample was applied on copper grids, preliminary negatively glow discharged using the EM ACE600 (LEICA microsystems), and incubated at room temperature for 1 min. The excess of the sample was blotted away and the grid was stained with 3 μL of 1% uranyl acetate for 20 s. The excess stain was blotted away and the grid was dried at room temperature. Grids were images on a JEOL 1400 microscope, operated at 120keV, equipped with a Orius 1000 GATAN camera.

##### 8.6 Immunogold electron microscopy

For Immuno-EM, nickel grids, coated with formvar film stabilised by 4 nm of continuous carbon, were negatively glow discharged using the EM ACE6000 (LEICA microsystems). 3 μL of the purified isolated conjugative pilus tips was applied on each grid before fixation with paraformaldehyde 1% diluted in buffer A (100 mM Tris [HCl], 200 mM NaCl, pH 8). Then, the grids were subjected to immunogold labelling as follows. The grids were blocked with blocking buffer (buffer A supplemented with SNC 10%, BSA 1%, BSAc 0.1%, pH 8) for 15 min two times, followed by incubation on a drop of either 1 : 1000 diluted primary anti-bodies anti-STREP (PA5-114453) with antibody-buffer (buffer A supplemented with SNC 1%, BSA 1%, BSAc 0.1%) or antibody-buffer only (for negative control) for 60 min at room temperature. Following four washes with the antibody-buffer, grids were incubated for 1 h with goat antirabbit Ig conjugated to 10 nm colloidal gold (BBI ; 1:30 diluted in Ab-buffer). Then, the grids were washed two times for 5 min with antibody-buffer followed by two times for 5 min with buffer A. Finally, the grids were fixed using 2.5% Glutaraldehyde diluted in buffer A for 5 min, washed four times for 3 min in H2O droplets and contrasted using Uranyl acetate 1% for 20 sec prior to examination.

#### 9. Experimental validation: Discovery of a novel F-box protein

The following procedures describe the HInt-based identification and experimental validation of Ydr249C as a novel F-box-like protein in *Saccharomyces cerevisiae*.

##### 9.1 Identification of a novel F-box protein using HInt

HInt was applied to the complete annotated *Saccharomyces cerevisiae* proteome (GenBank assembly GCF_000146045.2) using Skp1 (P52286) as the bait protein. Candidate proteins were filtered based on predicted subcellular localization, retaining only proteins predicted to localize to the cytoplasm or nucleus and lacking a signal peptide, thereby reducing the candidate set from 6033 to 4250 proteinsThe complete screening was completed in 193h 36min 59s. Among the resulting candidates, HInt identified Ydr249C (Q03787) as a putative Skp1-interacting protein (Supplementary Fig. 5a). Structural analysis of the predicted Ydr249C–Skp1 interaction model and interface residues revealed an interaction surface compatible with an F-box-like domain, providing additional support for Ydr249C as a candidate F-box protein.

##### 9.2 Yeast strains and plasmids

All yeast strains used in this study are isogenic derivatives of BY4741[43]. Cdc53-SmBiT, Ydr249C-LgBiT expressing strains were isolated from previously described haploid genome wide libraries[41]. The Δ(F-box) mutant of Ydr249C C-terminally fused to LgBiT was obtained by integrating a plasmid harbouring the corresponding constructs and a hygromycin resistance cassette into the endogenous *YDR249C* loci. The *skp1-3* strain was obtained from a collection of temperature-sensitive mutants provided by Charles Boone's laboratory. These strains were then crossed and sporulated using conventional methods to obtain haploid NanoBiT strains expressing Cdc53-SmBiT, Ydr249C-LgBiT (wild-type or Δ(F-box)) and wild-type or mutated Skp1 (Supplementary Fig. 5b).

##### 9.3 NanoBiT assays

NanoBiT yeast strains were cultured in SC(MSG) medium (1.7 g/L yeast nitrogen base without amino acids and ammonium sulfate, 2 g/L amino acid mix, 20 g/L glucose, 1g/L and monosodium glutamate) at 30°C. Overnight cultures were diluted twenty times in fresh SC(MSG) and further grown for 3 hours at 30°C. Optical density at 600 nm ($OD_{600}$) and luminescence measurements were performed for each culture in at least three technical replicates. For $OD_{600}$ measurements 125 µL of each culture were transferred to transparent 96-well plates (Falcon) and absorbance was recorded using a plate reader (Ensight, Perkin Elmer). For luminescence measurements 40 µL of each culture were transferred to white 96-well half-area microtiter plates (Greiner Bio-One) previously filled with 40 µL per well of SC(MSG) containing 100 µM furimazine. The microplates were shaken at 1200 RPM (2 mm amplitude) and incubated for 3 minutes before recording the luminescence signals with a luminometer (Ensight, Perkin Elmer). NanoBiT signals were computed as background subtracted luminescence, corrected for the $OD_{600}$ and normalised to 1 using the median of the three replicates of wild-type NanoBiT strains.

##### 9.4 SDS-PAGE and blotting

Protein extracts prepared from the different yeast strains were separated by SDS-PAGE (4-20% gradient) and blotted on nitrocellulose using standard protocols by chemiluminescence after addition of 1µM HiBiT peptide, to reconstitute an active luciferase, and 50 µM furimazine, as described[44].

### EXTENDED DATA FIGURES



a

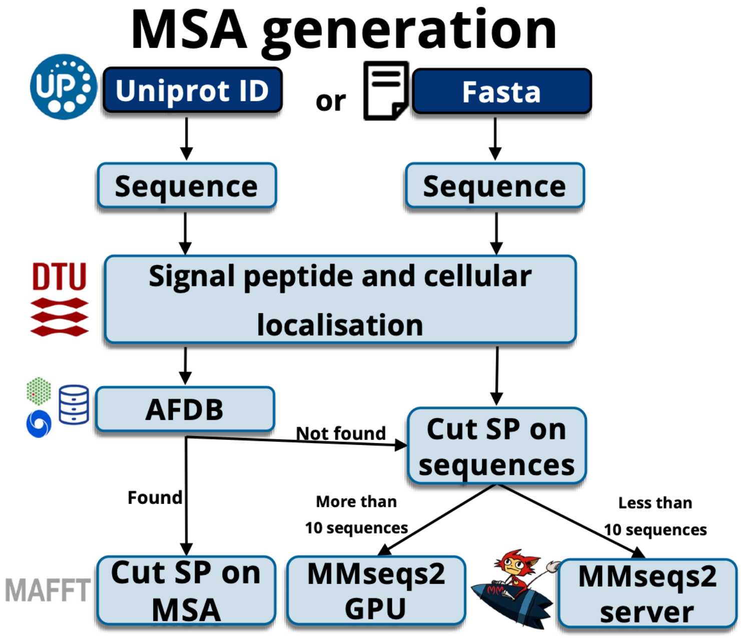


b

| | Number of MSA | Colab Fold MMseqs2 server | MMseqs2 GPU | HInt pipeline | Speed-up Server \| GPU |
|---|---|---|---|---|---|
| list 1 | 12 | 11min 26s | 4min 49s | 20.67s | **33x \| 14x** |
| list 2 | 34 | 19min 44s | 5min 31s | 36.46s | **32x \| 9x** |
| list 3 | 183 | 2h 41min 31s | 29min 28s | 6min 6s | **26x \| 5x** |
| list 4 | 785 | 11h 58min 1s | 2h 0min 2s | 15min 50s | **45x \| 8x** |
| list 5 | 6033 | 3d 19h 30min 5s* | 14h 22min 47s | 1h 38min 1s | **56x \| 14x** |

**Extended Data Fig. 1 | Accelerated MSA generation in HInt.**

**a, MSA generation workflow.**
The multiple sequence alignment (MSA) generation module follows a decision tree based on the input format (UniProt accession or FASTA sequence) and the availability of precomputed alignments. When a UniProt accession is provided, HInt first retrieves precomputed MSAs from the AlphaFold Protein Structure Database (AFDB). Otherwise, or when no precomputed alignment is available, MSAs are generated using GPU-accelerated MMseqs2. If fewer than 10 proteins require MSA generation, HInt automatically falls back to the MMseqs2 server instead of the GPU-accelerated MMseqs2 workflow. Signal peptide prediction is performed upstream, and predicted signal peptides are removed from the query sequence and corresponding MSA before downstream analyses.

**b, Benchmark of MSA generation.**
The table compares the execution time required to generate MSAs using the ColabFold MMseqs2 server (v1.5.5), GPU-accelerated MMseqs2 (v1.6.1), and the HInt workflow.

Benchmarks were performed on five datasets ranging from 12 to 6,033 proteins using 48 CPU cores and a single NVIDIA RTX 6000 Ada GPU. HInt achieved an average 47-fold speed-up over the ColabFold MMseqs2 server and an 11-fold speed-up over standalone GPU-accelerated MMseqs2. The value marked with an asterisk corresponds to an extrapolated runtime.

a

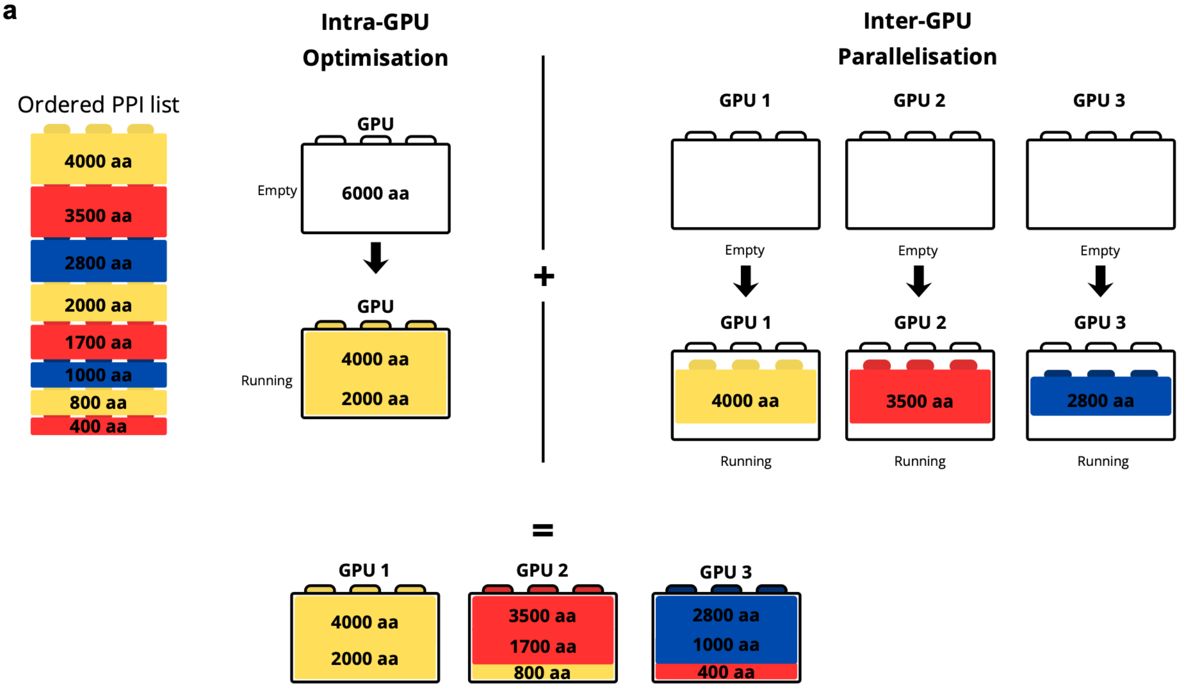


b

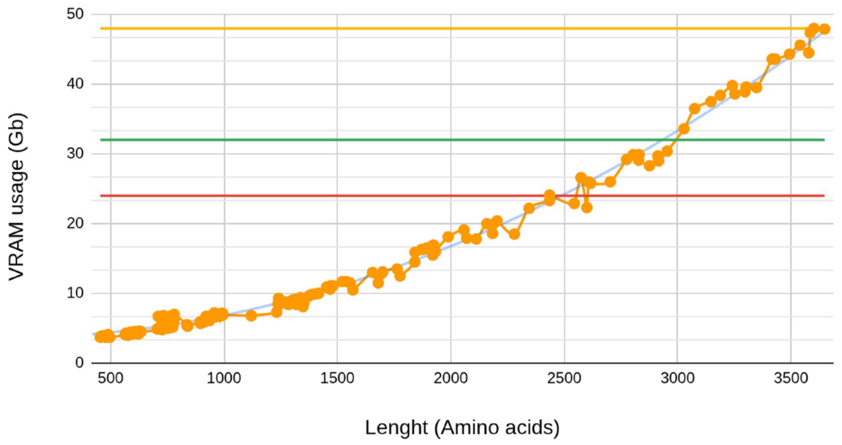


c

| | Number of PPIs | AF-M | HInt (AF-M) | **Speed-up** | AF3 | HInt (AF3) | **Speed-up** |
|---|---|---|---|---|---|---|---|
| list PPI 1 | 10 | 5h 13min | 1h 22min | **3.8x** | 42min 33s | 10min 56s | **3.9x** |
| list PPI 2 | 100 | 2d 9h 28min | 12h 27min | **4.6x** | 7h 45min | 1h 35min | **4.9x** |
| list PPI 3 | 1000 | 24d 16h 15min* | 5d 3h 33min | **4.8x** | 3d 6h 36min | 15h 45min | **5.0x** |

**Extended Data Fig. 2 | GPU scheduling and benchmarking of the HInt workflow.**

**a, GPU scheduling strategy implemented in HInt.**
HInt accelerates protein–protein interaction (PPI) prediction by combining two complementary scheduling strategies: intra-GPU memory-aware batching, which packs compatible prediction jobs onto the same GPU according to their estimated VRAM requirements, and inter-GPU parallelisation, which distributes prediction jobs across multiple GPUs. Each block represents

a predicted protein complex, with its size proportional to the combined sequence length of the interacting proteins. PPIs are first sorted by total sequence length and then dynamically assigned to maximize GPU utilisation while minimizing overall runtime.

**b**, **GPU memory requirements for AlphaFold-Multimer.**
GPU memory (VRAM) usage is shown as a function of the combined sequence length of the predicted protein complex. Measurements were obtained from 259 AlphaFold-Multimer predictions performed on NVIDIA RTX 4500 Ada (24 GB) and RTX 6000 Ada (48 GB) GPUs. Horizontal lines indicate the memory capacities of 24 GB, 32 GB, and 48 GB GPUs. The relationship between sequence length and VRAM usage was modeled using polynomial regression.

**c, Computational benchmark of the structure prediction stage.**
Execution times are compared for the structure prediction stage of three PPI datasets using the standard AlphaFold-Multimer and AlphaFold 3 workflows and the corresponding HInt implementations. Only 3D structure prediction time is included; MSA generation and interaction scoring are excluded. HInt achieves an approximately fivefold reduction in prediction time through GPU scheduling and memory-aware batching, independently of the underlying AlphaFold version. The average length of all PPI is 1222 amino acids. This benchmark was performed using four NVIDIA RTX 4500 Ada GPUs (24 GB).

**Extended**

ED Figure 3 Rouger et al. 2026

**a**

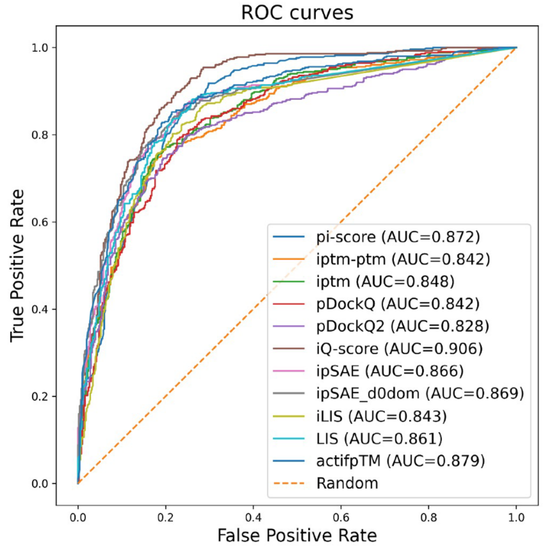


**b**

| | pi-score | ipTM-pTM | ipTM | pDockQ | pDockQ2 | iQ-score | ipSAE | ipSAE_d0dom | iLIS | LIS | actifpTM |
|---|---|---|---|---|---|---|---|---|---|---|---|
| **Thresholds** | 0.05 | 0.5 | 0.5 | 0.23 | 0.23 | 50 | 0.6 | 0.6 | 0.223 | 0.203 | 0.65 |
| **TP** | 461 | 366 | 372 | 407 | 207 | 368 | 234 | 351 | 409 | 410 | 435 |
| **FN** | 39 | 134 | 128 | 93 | 293 | 132 | 266 | 149 | 91 | 90 | 65 |
| **FP** | 159 | 86 | 87 | 131 | 29 | 57 | 29 | 60 | 118 | 102 | 124 |
| **TN** | 341 | 414 | 413 | 369 | 471 | 443 | 471 | 440 | 382 | 398 | 376 |
| **Sensitivity** | 0.922 | 0.732 | 0.744 | 0.814 | 0.414 | 0.736 | 0.468 | 0.702 | 0.818 | 0.82 | 0.87 |
| **Specificity** | 0.682 | 0.828 | 0.826 | 0.738 | 0.942 | 0.886 | 0.942 | 0.88 | 0.764 | 0.796 | 0.752 |
| **PPV** | 0.744 | 0.810 | 0.810 | 0.757 | 0.877 | 0.866 | 0.890 | 0.854 | 0.776 | 0.801 | 0.778 |
| **NPV** | 0.897 | 0.755 | 0.763 | 0.799 | 0.616 | 0.770 | 0.639 | 0.747 | 0.808 | 0.816 | 0.853 |

**Data Fig. 3 | Benchmark of protein-protein interaction scoring metrics.**

**a**, **ROC analysis of protein-protein interaction scoring metrics.**
Receiver operating characteristic (ROC) curves comparing eleven scoring metrics for protein–protein interaction evaluation. All methods were benchmarked using the same dataset comprising 500 experimentally validated positive interactions from the IntAct database and 500 negative interactions from the Negatome database. Protein complexes were predicted using AlphaFold-Multimer, and each scoring metric was evaluated on the same set of predicted structures. The area under the ROC curve (AUC) is indicated for each method.

**b**, **Performance summary of protein–protein interaction scoring metrics.**
Performance metrics calculated at the selected threshold for each scoring method. The table reports the optimal threshold, true positives (TP), false negatives (FN), false positives (FP), true negatives (TN), sensitivity, specificity, positive predictive value (PPV), and negative predictive value (NPV).

**a**

| Uniprot | Gene | DeepLoc | Signal peptide | iQ-score vs P33790 |
|---|---|---|---|---|
| P04737 | traA | Extracellular\|Cytoplasmic Membrane | Yes | 63.04 |
| P15069 | traH | Outer Membrane\|Periplasmic | Yes | 58.72 |
| P47737 | yubQ | Extracellular\|Outer Membrane\|Periplasmic | Yes | 28.04 |
| P18035 | trbB | Periplasmic | Yes | 26.47 |
| Q9JMT7 | yuaC | Extracellular\|Outer Membrane | Yes | 25.87 |
| P41066 | traK | Periplasmic | Yes | 14.28 |
| P34211 | yuaR | Extracellular\|Cytoplasmic | Yes | 0 |
| Q9JMS2 | yuaS | Extracellular\|Cytoplasmic | Yes | 0 |
| P08868 | yuaZ | Extracellular\|Cytoplasmic\|Outer Membrane | Yes | 0 |
| Q9JMR4 | yubK | Extracellular\|Cytoplasmic Membrane | Yes | 0 |
| P18472 | traW | Periplasmic | Yes | 0 |
| P18471 | traU | Extracellular\|Periplasmic | Yes | 0 |
| P18473 | trbC | Periplasmic | Yes | 0 |
| P24082 | traN | Extracellular\|Outer Membrane | Yes | 0 |
| A0A649URK2 | ybgA | Extracellular\|Cytoplasmic | Yes | 0 |
| Q84A26 | traF | Periplasmic | Yes | 0 |

**Extended Data Fig. 4 | HInt screening identifies the F-plasmid VirB5 homolog.**

a, **HInt screening results of the F-plasmid proteome using $TraG_{B6}$ as bait.**
Summary of HInt predictions obtained by screening $TraG_{B6}$ (P33790) against the F-plasmid proteome (110 proteins). The search incorporated biologically informed filters requiring a predicted periplasmic or extracellular localisation together with the presence of a signal peptide. The complete screen required 1h 52min and identified P15069 (TraH), highlighted in red, as the top candidate corresponding to the previously unidentified VirB5 homolog. The known interaction partner $TraA_{B2}$ (P04737), which interacts with $TraG_{B6}$ through a distinct interface, was also recovered, providing an internal positive control for the screen. The average combined sequence length of the predicted protein pairs was 948 amino acids. The screening was performed using 64 CPU cores and four NVIDIA RTX 4500 Ada GPUs.

**a**

| Name | Gene | DeepLoc | Signal peptide | iQ-score vs P52286 | InterPro entries |
|---|---|---|---|---|---|
| Q08496 | DIA2 | Nucleus | No | 83.68 | IPR001810, IPR036047 |
| Q03787 | YDR249C | Nucleus | No | 81.98 | |
| P07834 | CDC4 | Nucleus | No | 81.67 | IPR001810, IPR036047 |
| Q03899 | YDR131C | Cytoplasm\|Nucleus | No | 80.7 | IPR001810 |
| P47005 | DAS1 | Cytoplasm\|Nucleus | No | 80.5 | IPR001810, IPR036047 |
| Q05930 | MDM30 | Cytoplasm\|Nucleus | No | 80.44 | IPR001810, IPR036047 |
| Q12347 | HRT3 | Cytoplasm\|Nucleus | No | 80.25 | IPR036047 |
| Q05947 | UCC1 | Cytoplasm\|Nucleus\|Mitochondrion | No | 80.19 | |
| P38352 | SAF1 | Cytoplasm\|Nucleus | No | 80.15 | IPR036047 |
| Q04847 | ROY1 | Cytoplasm\|Nucleus\|Mitochondrion | No | 79.42 | IPR001810, IPR036047 |
| P39531 | RCY1 | Cytoplasm\|Golgi apparatus | No | 78.4 | |
| Q06640 | YDR306C | Cytoplasm\|Nucleus | No | 77.8 | |
| Q04511 | UFO1 | Cytoplasm\|Nucleus | No | 77.35 | IPR001810, IPR036047 |
| P38285 | AMN1 | Cytoplasm\|Nucleus | No | 77.26 | |
| P24814 | GRR1 | Cytoplasm\|Nucleus | No | 76.83 | IPR001810, IPR036047 |
| P42843 | SKP2 | Cytoplasm\|Nucleus\|Mitochondrion | No | 75.82 | IPR001810, IPR036047 |
| P39014 | MET30 | Nucleus\|Mitochondrion | No | 74.22 | IPR001810, IPR036047 |
| P38308 | COS111 | Cytoplasm\|Nucleus | No | 73.77 | IPR036047 |
| P0C5N4 | YHR073C-B | Cytoplasm\|Nucleus\|Mitochondrion | No | 73.41 | |
| Q06479 | YLR352W | Cytoplasm\|Nucleus | No | 73.09 | |
| P35203 | CTF13 | Cytoplasm\|Nucleus | No | 72.3 | |
| Q04922 | MFB1 | Cytoplasm | No | 71.34 | IPR001810, IPR036047 |

**b**

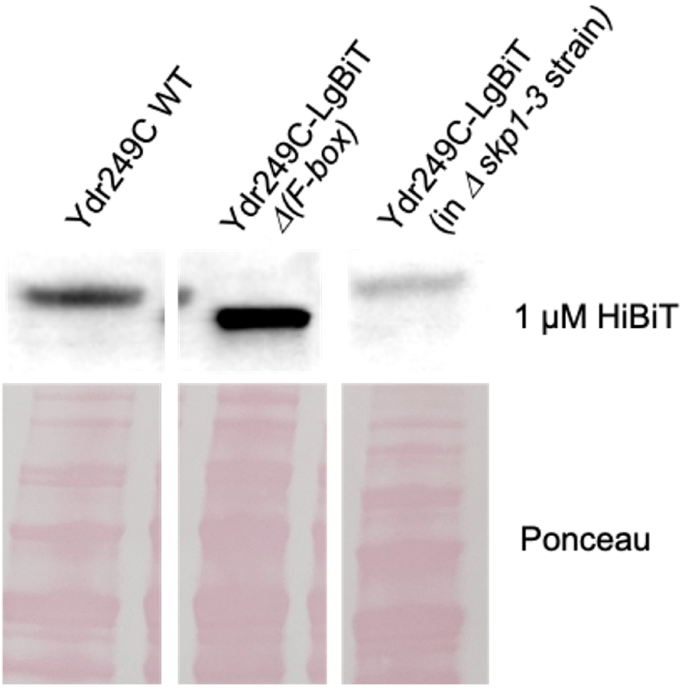


**Extended Data Fig. 5 | Genome-wide HInt screening identifies a previously unannotated F-box protein.**

**a, HInt screening of the *S. cerevisiae* proteome using Skp1 as bait.**
Summary of HInt predictions obtained by screening Skp1 (P52286) against the complete *Saccharomyces cerevisiae* proteome (6,033 proteins). The search incorporated biologically informed filters requiring predicted nuclear or cytoplasmic localisation. The complete

screening was completed in 193h 36min 59s. Only proteins with an iQ-score greater than 70 are shown. Previously characterized F-box proteins are highlighted in green, whereas the previously unannotated candidate Ydr249C is highlighted in red. InterPro domains identified in each protein are indicated. The average combined sequence length of the predicted protein pairs was 682 amino acids. The screening was performed using 64 CPU cores and four NVIDIA RTX 4500 Ada GPUs.

**b, Expression of Ydr249C constructs used for NanoBiT assays.**
Blot confirming expression of the wild-type and Δ(F-box) Ydr249C-LgBiT constructs used in the different NanoBiT strain. Ydr249C-LgBiT protein levels are reduced in the *skp1-3* background, suggesting that Skp1 may influence Ydr249C abundance or stability. Ponceau staining serves as a loading control.
